# Natural selection does not affect the estimates of effective population size based on linkage disequilibrium

**DOI:** 10.1101/2021.08.16.456457

**Authors:** Irene Novo, Enrique Santiago, Armando Caballero

**Author notes:** Correspondence (I.N.).

## Abstract

The effective population size (*N_e_*) is a key parameter to quantify the magnitude of genetic drift and inbreeding, with important implications in human evolution. The increasing availability of high-density genetic markers allows the estimation of historical changes in *N_e_* across time using measures of genome diversity or linkage disequilibrium between markers. Selection is expected to reduce diversity and *N_e_*, and this reduction is modulated by the heterogeneity of the genome in terms of recombination rate. Here we investigate by computer simulations the consequences of selection (both positive and negative) and of recombination rate heterogeneity in the estimation of historical *N_e_*. We also investigate the relationship between diversity parameters and *N_e_* across the different regions of the genome using human marker data. We show that the estimates of historical *N_e_* obtained from linkage disequilibrium between markers (*N*_*e*LD_) are virtually unaffected by selection. In contrast, those estimates obtained by coalescence mutation-recombination-based methods can be strongly affected by it, what could have important consequences for the estimation of human demography. The simulation results are supported by the analysis of human data. The estimates of *N*_*e*LD_ obtained for particular genomic regions do not correlate with recombination rate, nucleotide diversity, polymorphism, background selection statistic, minor allele frequency of SNPs, loss of function and missense variants and gene density. This suggests that *N*_*e*LD_ measures are merely indicative of demographic changes in population size across generations.

**Author summary:** The inference of the demographic history of populations is of great relevance in evolutionary biology. This inference can be made from genomic data using coalescence methods or linkage disequilibrium methods. However, the assessment of these methods is usually made assuming neutrality (absence of selection). Here we show by computer simulations and analyses of human data that the estimates of historical effective population size obtained from linkage disequilibrium between markers are unaffected by natural selection, either positive or negative. In contrast, estimates obtained by coalescence mutation-recombination-based methods can be strongly affected by it, which could have important consequences for recent estimations of human demography. Thus, only linkage disequilibrium methods appear to provide unbiased estimates of the population census size.

## Introduction

The effective population size (*N_e_*) is a parameter of paramount relevance in evolutionary biology, plant and animal breeding and conservation genetics, because its magnitude reflects the amount of genetic drift and inbreeding occurring in the population^1^. The effective size of a population depends on its demographic history and structure as well as the selection regime affecting the population^2–4^. Estimates of *N_e_* can be obtained by methods using information from genetic markers^3,5,6^, and those based on linkage disequilibrium (LD) between them are generally acknowledged to be reliable and robust.^7–8^ The idea behind these methods is that, for neutral loci in an isolated population LD is inversely proportional to the genetic distance (or recombination rate, *c*) between any pair of markers and the effective size of the population.^9^

With the increasing availability of high-density marker information, such as that of single nucleotide polymorphisms (SNP) panels and whole genome sequences for more and more species^10^, methods based on LD that allow an estimate of the temporal changes of *N_e_* in the recent past have been developed.^11–13^ The basic idea is that LD between pairs of SNPs at different genetic distances provides differential information on *N_e_* at different time points in the past. Thus, Hayes and colleagues^11^ suggested that LD between loci with a recombination rate *c* approximately reflects the ancestral effective population size 1/(2*c*) generations ago. Thus, they proposed to estimate *N_e_* at a given generation *t* from pairs of SNPs at a genetic distance 1/(2*t*) Morgans. This method has become increasingly popular for estimating the past and present *N_e_* in human^12,14^ and livestock^15,16^ populations, and a number of bioinformatic tools have been developed to allow its implementation (e.g. Hollenbeck et al.^17^).

The original application of the above method for estimating historical *N_e_* is, however, restricted to the assumption of constant or linear population growth or decline.^11^ Thus, drastic population size changes such as bottlenecks or sudden severe declines in census size, which are common in natural populations or at the start of breeding programs, cannot be detected accurately with this method. A late development has been shown to accurately detect drastic changes in historical *N_e_*.^13^ Over relatively recent timespans of about 200 generations back in time, the method has been shown to be more accurate than other alternative coalescence and mutation-recombination-based methods, such as MSMC^18^ and Relate,^19^ which are expected to be applied for long term evolutionary estimations.

All methods of estimation of past *N_e_* trajectories assume neutrality (absence of selection). It is well known, however, that selection significantly reduces *N_e_*,^2,20–21^ particularly in regions of low recombination.^2,22–27^ Natural populations are predicted to encode many deleterious variants^28–29^ that can affect the outcome of *N_e_* estimation methods. The fate of these variants, as well as of that of advantageous ones, also relies on linkage, because selection is less effective in genomic regions of low recombination.^30^

Genomes are also heterogeneous for genetic variation due to differences in recombination rates across chromosomal regions^31–34^ and because of the differential impact of natural selection on them.^35^ For example, selective sweeps of favourable mutations are expected to hitch-hike close-by neutral SNPs producing sharp decreases in diversity in linked regions.^36–38^ Ignoring the heterogeneity in *N_e_* may lead to biased estimates of past demography^39^ and, in fact, a heterogeneity of *N_e_* across the genome has been found for different eukaryotic species.^40–42^

It has been shown that selective sweeps of favourable mutations generate LD between close-by neutral loci,^43^ although this LD increase can be negligible^44^ and, in any case, it is transient and disappears quickly.^38,45^ If so, estimates of *N_e_* based on LD would be unaffected by selection. In this paper we assess the impact of selection on the estimation of historical *N_e_* using the method of Santiago and colleagues^13^, which is based on linkage disequilibrium between SNP markers, in comparison with other coalescence mutation-based methods, MSMC^18^ and Relate.^19^ Using individual-based forward simulations, we compare the estimates of historical *N_e_* provided by these methods assuming selective sweeps of favourable mutations and background selection on deleterious mutations, and considering the heterogeneity in recombination rates across the genome. In addition, we investigate the relationship between the estimates of linkage disequilibrium *N_e_* and other diversity and genomic parameters across the human genome, using SNP data obtained from genome sequencing of Finnish^46^ and Koryaks^47^ populations. We obtained the correlation between the estimates of local *N_e_* and several diversity variables over small windows across the genome. Both the simulation results and the analyses of human genome data provide strong evidence that estimates of effective population size based on LD reflect the true census of reproductive individuals in populations.

## Results

### Effect of selection and recombination on the estimation of *N_e_*

Figure 1 shows the joint effect of recombination and selection on the estimates of *N_e_*, for a population of constant size of *N* =1,000 individuals assuming different recombination rates (*RR*) per Mb across the genome. Under a neutral scenario, linkage disequilibrium estimates (*N_eLD_*) provide virtually unbiased estimates of *N* for all recombination rates. The same result can be observed, as expected, for estimates of *N_e_* based on nucleotide diversity (*π*) and calculated as *N_eπ_* = *π*/(4*μ*), where *μ* is the nucleotide mutation rate, assumed also to be constant across the genome. Estimates obtained from Relate (*N_eRelate_*) and MSMC (*N_eMSMC_*) give also precise estimates of *N* except for the most extreme cases of recombination rate 5 and 0.01 cM/Mb.

**Figure 1.**
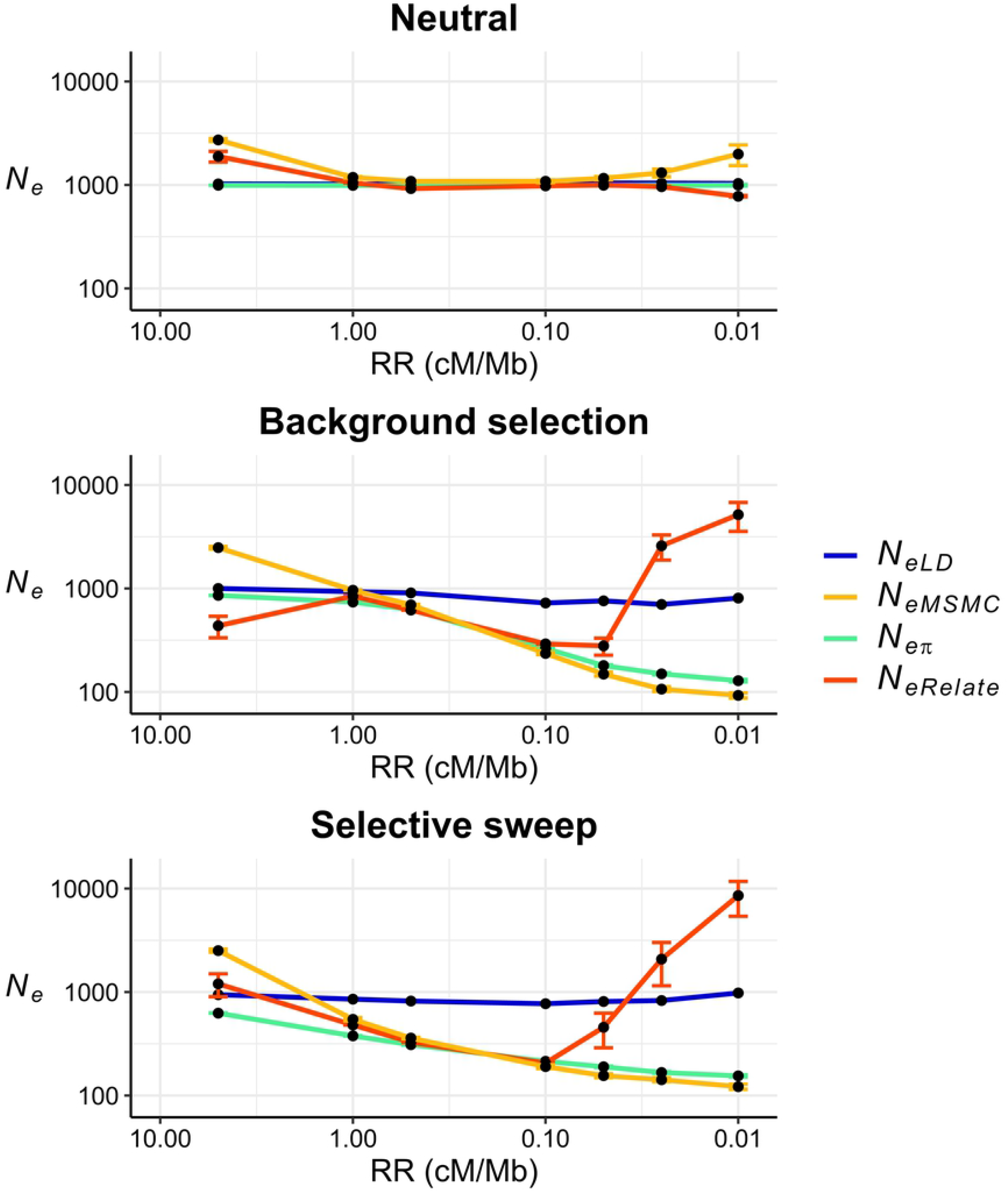
Estimates of effective population size from linkage disequilibrium (*N_eLD_*, sample size of *n* = 100 individuals), Relate (*N_eRelate_*, *n* = 100), MSMC (*N_eMSMC_*, *n* = 4), and from nucleotide diversity (*π*), this latter calculated as *N_eπ_* = *π*/(4*μ*), where *μ* is the nucleotide mutation rate, for scenarios with different recombination rates (*RR* in cM/Mb) uniform across the genome. Simulations assume a constant population size of *N* = 1,000 individuals under neutrality, background selection and selective sweeps, with constant mutation rate *μ* = 10^-8^ per base per generation. Estimates were obtained including windows of recombination rate between pairs of SNPs ranging from *c* = 0.0025 to 0.0250 for *N_eLD_* and averaging historical estimates of *N_e_* between generations 150 to 350 for *N_eRelate_* and *N_eMSMC_*. Error bars represent one standard error above and below the mean of the simulation replicates. *N_eLD_* does not have errors as single estimates were obtained using all pairs of SNPs from all replicates. All simulations were run for up to 100 replicates.

Under background selection and selective sweep models, *N_eLD_* estimates give the same results as for the neutral model, indicating that these estimates are very little or not affected by selection, either negative or positive (Fig. 1). As expected, estimates of *N_eπ_* diverges from the true census size (*N*) of the population, with decreasing values as recombination rate decreases. Relate and MSMC estimates show a pattern similar to that of *N_eπ_* but *N_eRelate_* estimates increase sharply for tight linkage scenarios.

### Simulation results for historical *N_e_* estimates

Figure 2 shows estimates of historical *N_e_* assuming a constant population census size (*N* = 1,000 or 10,000), either with a constant or a variable recombination rate. Estimates of historical *N_eLD_* reflect the population census size regardless of selection and the variability in genomic recombination rates. Estimates from *N_eRelate_* and *N_eMSMC_* can give unbiased values of *N* under a neutral and background selection model if the initial generations are discarded and population size is not too large (*N* = 1000). Otherwise, they can show important differences from the population size, particularly under a selective sweep model, where *N* is underestimated by *N_eRelate_*, and can be either underestimated or overestimated by *N_eMSMC_*. In general, variation in the recombination rate across the genome has a limited effect on the estimates of *N* with respect to a constant recombination rate model.

**Figure 2.**
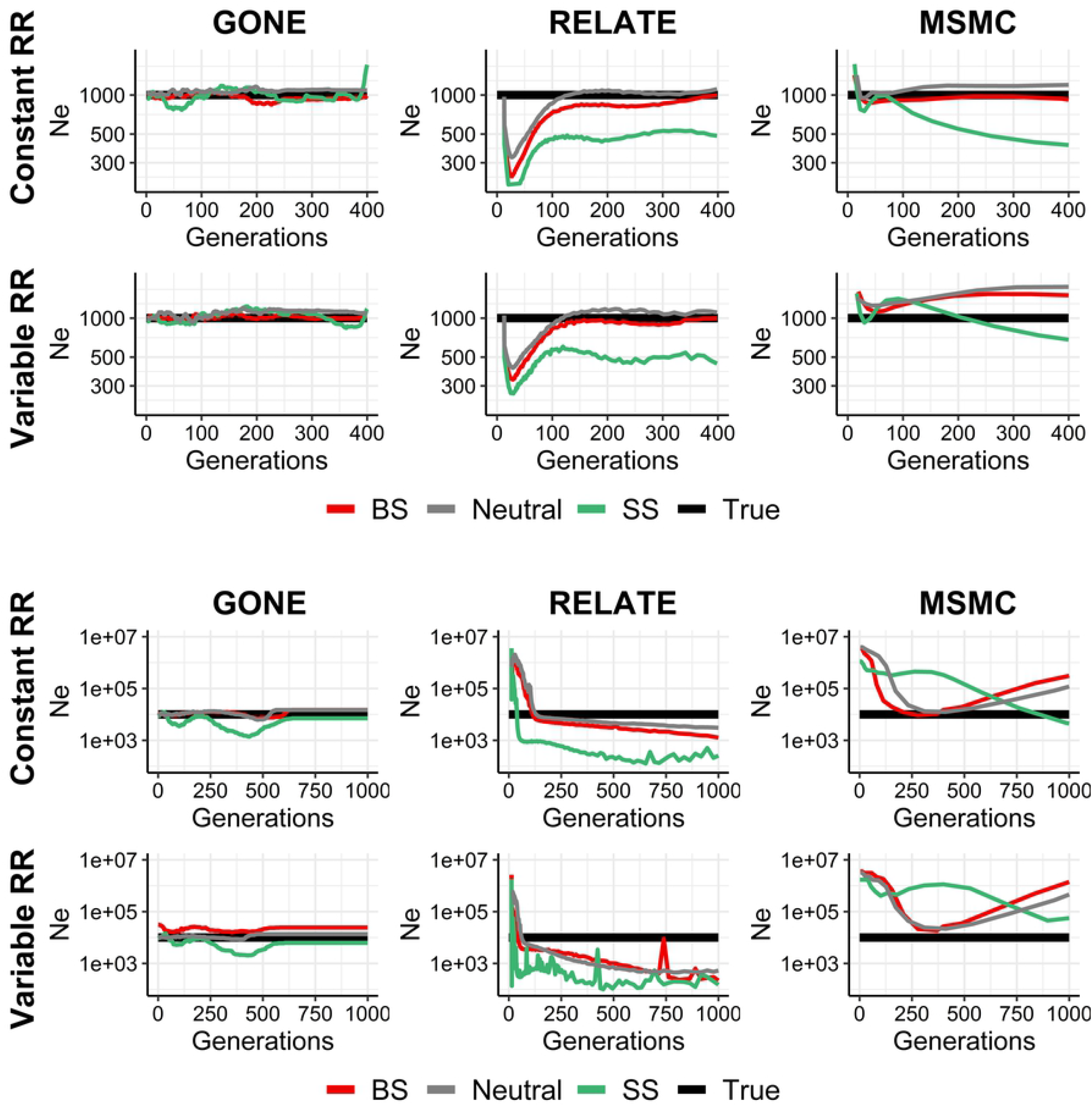
Estimates of historical *N_e_* with GONE (sample size *n* = 100 individuals), Relate (*n* = 100) and MSMC (*n* = 4) from the present generation (generation 0) back to 400 or 1,000 generations in the past. 100 replicates were run of simulations with constant population size (*N*) under neutrality (grey), background selection (BS, red) and selective sweeps (SS, green), with constant (1 cM/Mb) or variable recombination rates (*RR*), and constant mutation rate *μ* = 10^-8^ or 10^-9^ mutations per base per generation for *N* = 1,000 or *N* = 10,000, respectively. The true simulated *N* is shown in black.

Regarding historical estimates with variable *N_e_* (Fig. 3), estimates of *N_eLD_* predict accurately a recent demographic change in population size (occurred 30 generations in the past) regardless of selection and recombination rate heterogeneity. Relate’s estimates show a certain noise, particularly for selective sweeps and/or variable recombination rate scenarios, but give the approximate *N* value except for an underestimation in several selective sweep cases. MSMC’s estimates are also accurate, except for the most recent generations in the exponential *N* growth case. Selective sweeps tend to lead to an underestimation of *N_eMSMC_*, and recombination rate variability only biases estimates in the recent exponential growth case.

**Figure 3.**
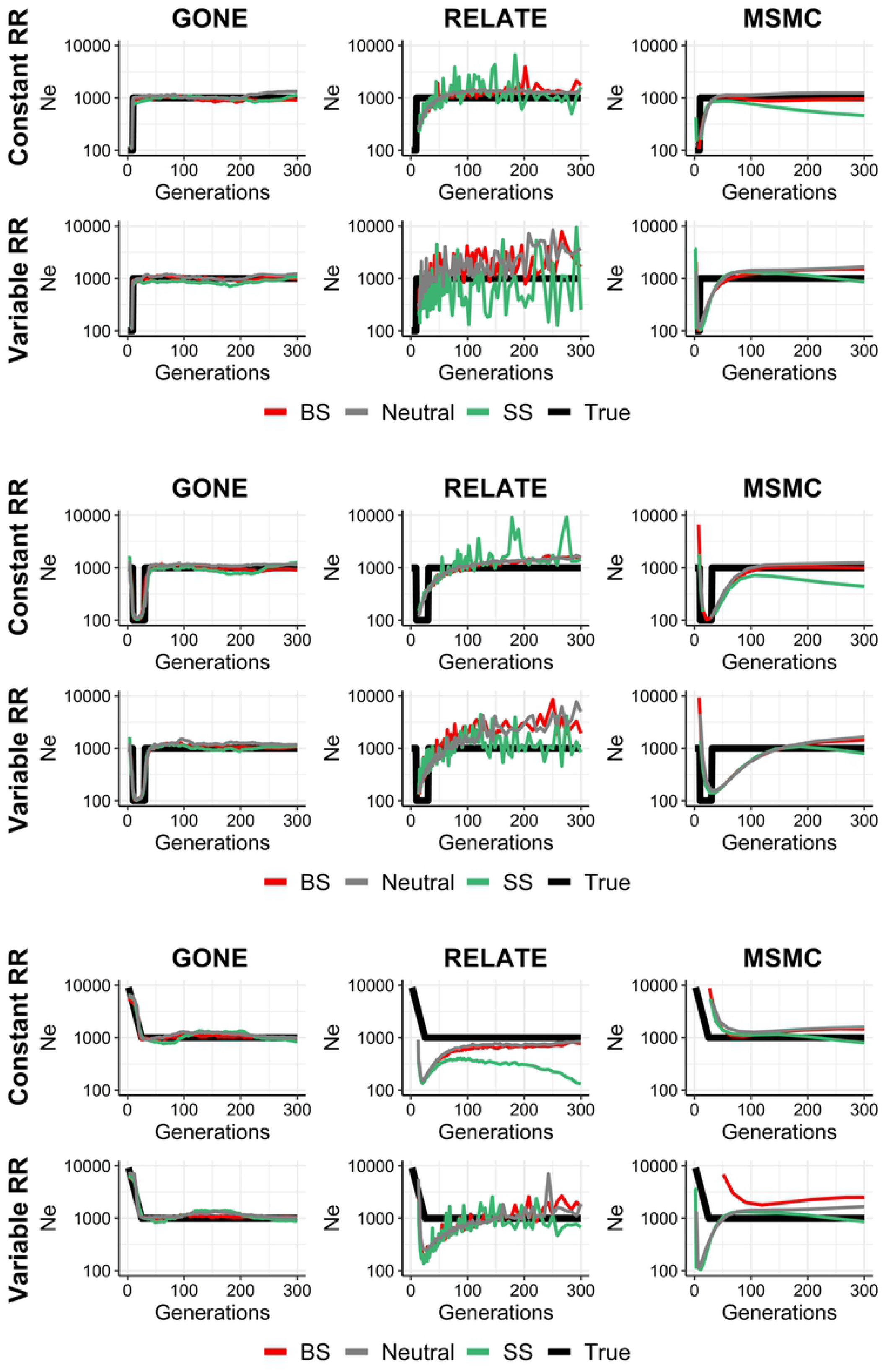
Estimates of historical *N_e_* with GONE (sample size *n* = 100 individuals), Relate (*n* = 100) and MSMC (*n* = 4) from the present generation (generation 0) back to 300 generations in the past. 100 replicates were run of simulations with constant population size (*N*) under neutrality (grey), background selection (BS, red) and selective sweeps (SS, green), with constant (1 cM/Mb) or variable recombination rate (*RR*), and constant mutation rate *μ* = 10^-8^ or 10^-9^ mutations per base per generation for *N* = 1,000 or *N* = 10,000, respectively. The true simulated *N* is shown in black.

When population size changes occur in more ancient times (around 300 generations ago; Fig. 4), both *N_eRelate_* and *N_eMSMC_* are unable to detect these demographic changes, while *N_eLD_* is generally more accurate. Nevertheless, *N_eLD_* estimates can be noisy, and sometimes, unable to detect the change in *N* when selection is present. A variable recombination rate does not substantially affect the estimates.

**Figure 4.**
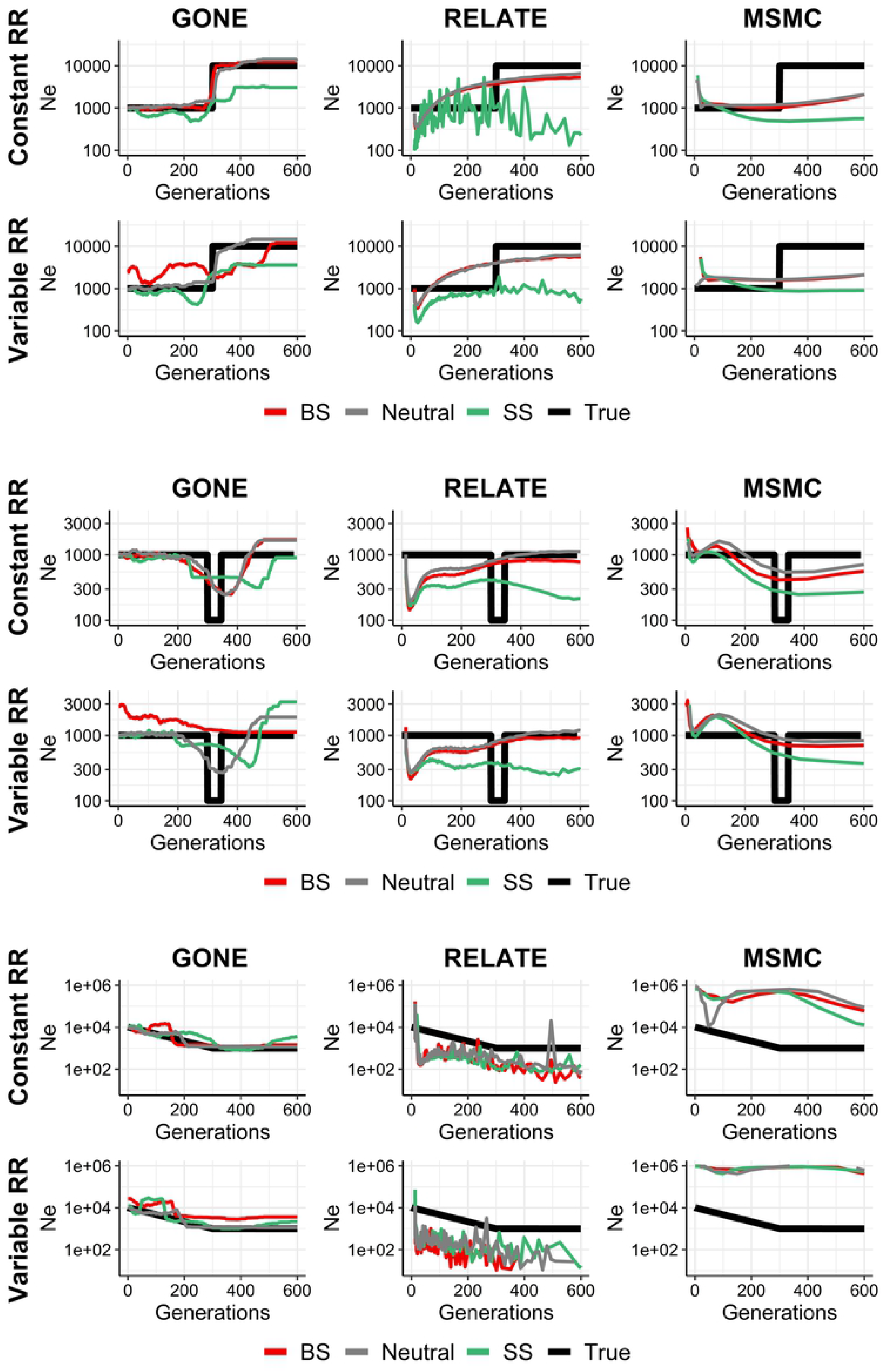
Estimates of historical *N_e_* with GONE (sample size *n* = 100 individuals), Relate (*n* = 100) and MSMC (*n* = 4) from the present generation (generation 0) back to 600 generations in the past. 100 replicates were run of simulations with constant population size (*N*) under neutrality (grey), background selection (BS, red) and selective sweeps (SS, green), with constant (1 cM/Mb) or variable recombination rate (*RR*), and constant mutation rate *μ* = 10^-8^ or 10^-9^ mutations per base per generation for *N* = 1,000 or *N* = 10,000, respectively. The true simulated *N* is shown in black.

### Correlation between regional estimates of *π* and *N_eLD_* with other diversity parameters using human data

The correlation between the average nucleotide diversity (*π*) in each genomic region and the other related variables followed the expected trends (Fig. 5a). A strong positive correlation was found between *π* and recombination rate *RR*, the background selection statistic *B*, nucleotide polymorphism *P* and MAF of SNPs. Nucleotide diversity was also weakly negatively correlated with loss-of-function, missense mutations and gene density, but only significantly for the Finnish population. Finally, no correlation was found between *π* and *N_eLD_*.

**Figure 5.**
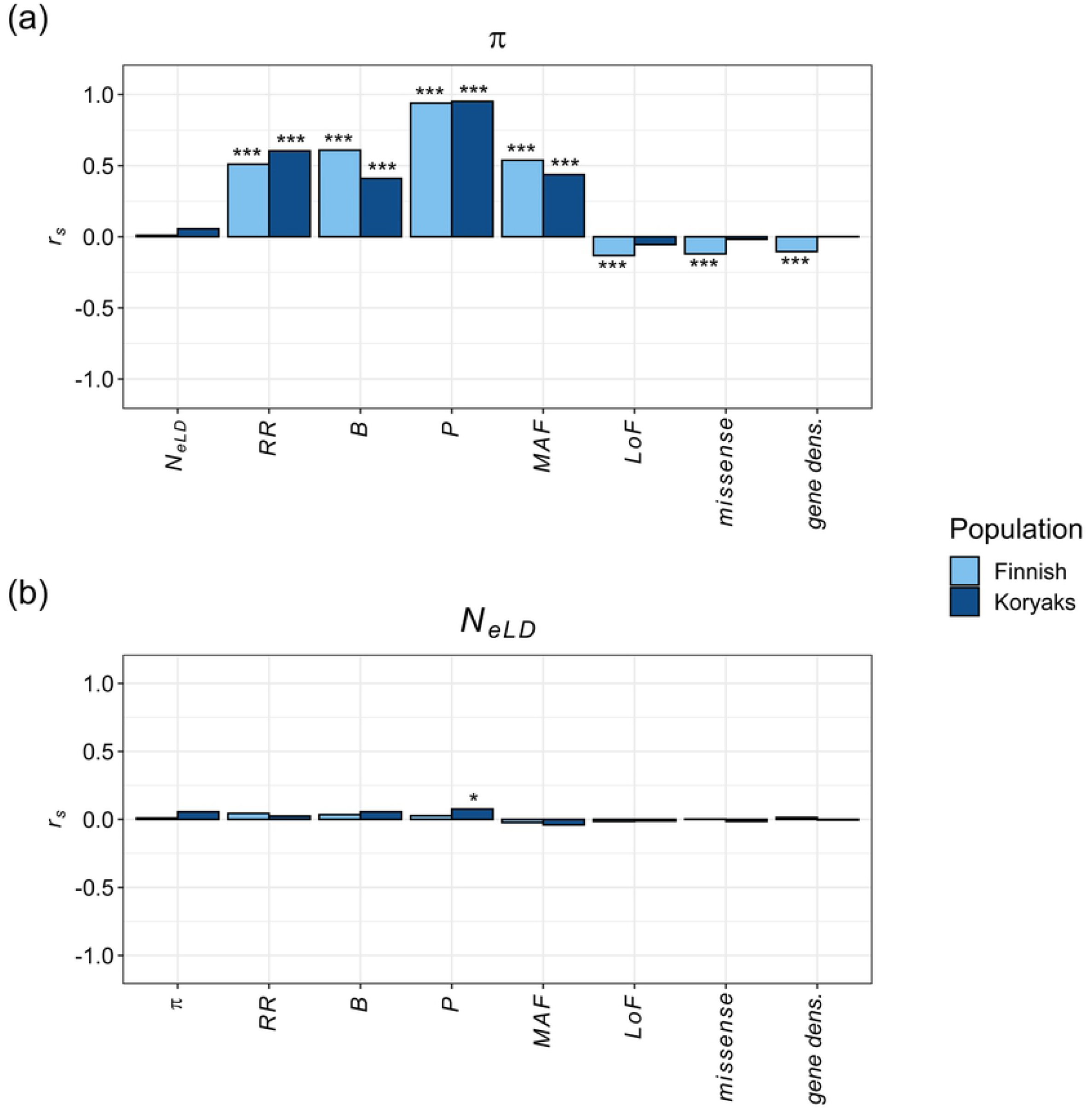
Spearman’s correlation coefficient (*r*) between nucleotide diversity (*π*, panel a) or estimates of linkage disequilibrium effective population size (*N_eLD_*; panel b) and different diversity variables, estimated within genomic regions. *RR*: recombination rate; *B*: B statistic; *P*: nucleotide polymorphism; MAF: Minor Allele Frequency; *LoF*: number of Loss of Function variants; *missense*: number of missense variants; *gene dens*: gene density. Estimates are based on samples of *n* = 99 for Finnish and *n* = 16 for Koryaks populations. P-values: * < 0.05, *** <0.001.

The correlations among all genetic variables studied are shown in the Supplemental material (Fig. S2), and follow the expected trends. For example, recombination rate *RR*, the *B* statistic, polymorphism *P* and MAF of SNPs were highly positively correlated among them. The *B* statistic was strongly negatively correlated with gene density and deleterious variation (LoF, missense), and these latter were highly correlated among them.

The mean estimates (and standard deviation) of *N_eLD_* across regions were 8,377 (5,462) for the Finnish population and 2,034 (3,660) for the Koryaks population. Thus, the standard deviation of the regional estimates of *N_eLD_* relative to a mean of one, for comparison, were 0.65 for the Finnish population and 1.80 for the Koryaks population.

The correlation between the estimates of *N_eLD_* and other diversity parameters for different genomic regions are shown in Figure 5b. The correlations did not follow the trends observed for nucleotide diversity. In particular, there was no significant correlation between *N_eLD_* and *π*, *RR*, *B*, *P*, MAF, loss-of-function variants, missense variants and density of genes, except for a small significant correlation between *N_eLD_* and *P* for the Koryaks population.

## Discussion

Our results show that the estimates of census population size obtained from linkage disequilibrium between pairs of SNPs (*N_eLD_*) are virtually unaffected by either positive or negative selections. This has been deduced from simulation data assuming constant or variable population size and different selection models (Figs. 1–4). The simulation results are supported by those obtained from real human genomic data. The estimates of *N_eLD_* in genomic windows are uncorrelated with recombination rate, the *B* statistic (which quantifies the strength of background selection), nucleotide diversity and polymorphism, as well as the number of deleterious variants (loss-of-function and missense variants) and density of genes (Fig. 5). Thus, the results show that the estimates of *N_eLD_* are basically unaffected by selection.

Although positive selection is known to generate linkage disequilibrium between neutral loci closed to selective loci,^43^ this effect is transient and disappears very fast.^38,45^ Stephan and colleagues^45^ showed that the increase in linkage disequilibrium between two neutral loci occurs if both are at less than a genetic distance of *c* = 0.1*s*, where *s* is the selection coefficient of the advantageous allele in homozygosis. We performed simulations with *s* = 0.02, which implies that LD is generated between loci located at a genetic distance of *c* = 0.002, or 0.2 cM. In the most extreme linkage scenario simulated in Figure 1 we assumed a rate of recombination of *RR* = 0.01 cM per Mb, which implies a total genetic distance of 1 cM for the whole simulated genome sequence of 100 Mb. Even in this extreme scenario, the estimates of *N_eLD_* are unaffected by positive selection. Note that, although some transient disequilibrium may be generated during selective sweeps for close-by neutral loci, the estimates of *N_eLD_* are based on all pairs of loci across the genome, so it is expected that many pairs are not affected by any effect of selection on linkage.

In contrast, and as it is well known (e.g. Charlesworth^2^, Santiago and Caballero^27^ Gillespie^48^), the nucleotide diversity effective population size (*N_eπ_*) is drastically reduced in regions of low recombination under selection (Fig. 1). The observed strong correlations between the regional genomic values of *π* and the recombination rate, the background selection statistic, and the deleterious variants from human data (Fig. 5a), also confirms this observation. Estimates of *N_e_* obtained by mutation-recombination-based coalescence methods (MSMC and Relate) are also affected by selection (Figs. 1–4). In fact, a Relate Selection Test has been proposed based on estimating the speed of spread of a particular lineage relative to other competing lineages.^19^ MSMC estimates generally follow the pattern of *N_eπ_* estimates except for large recombination rates, for which it provides overestimates of the true population size (Fig. 1). Relate estimates also follow *N_eπ_* estimates down to recombination rates of 0.1 cM/Mb but, for lower values, the estimates increase drastically above the true population size (Fig. 1). These coalescence methods have been used to investigate ancient demography of human populations and are not generally applicable to short-term historical changes in population size (see Figs. 2–4). In fact, it has been acknowledged that MSMC with 8 haplotypes works for estimations before about 70 generations in the past, i.e. about 2,000 years for human populations^18^, and Relate seems to discriminate after about 1,000 years back.^19^ Thus, MSMC was able to detect the out-of-Africa bottleneck in non-African populations from 200,000 years ago until 50,000 years ago,^18^ while Relate detected it from 40,000 to 20,000 years ago.^19^ The possible impact of selection on these demographic inferences is an issue to be further investigated.

The effect of recombination rate heterogeneity on historical *N_e_* estimates is not very noticeable in most cases (Fig. 2–4), particularly when GONE and MSMC are used. This is in accordance with the analyses made by Schiffels and Durbin (their Supplementary Figure 4)^18^, showing that simulated estimates from MSMC obtained using chunks of the human recombination map do not differ much from those using a constant recombination rate. For GONE estimates, varying recombination rate seems to only affect some of the background selection scenarios, while estimates obtained under a neutral and selective sweep models remain similar to those obtained with the constant recombination rate scenario. For Relate estimates, recombination rate heterogeneity seems to affect the estimates of recent demographic changes (Figure 3), generating noisier estimations. MSMC estimates are, in general, only slightly affected by variable recombination rate.

Gossmann and colleagues^40^ quantified the heterogeneity of *N_e_* across the genome of ten eukaryotic species (including humans) through the nucleotide diversity of genome sites and accounting for the differences in mutation rate between loci by considering the divergence between species. Thus, they obtained estimates of *N_eπ_*, finding a modest but statistically significant variability of *N_e_* for most species. Gossmann and colleagues^40^ found that *N_e_* was only positively correlated with recombination rate for *Drosophila* (*r* = 0.45), and negatively correlated with gene density for *Arabidopsis* (*r* = −0.11) and humans (*r* = −0.19). These correlations generally agree with those found for nucleotide diversity in our analysis (Fig. 5a), i.e., *r* = 0.51 and 0.60 between *π* and *RR*, and *r* = −0.10 and 0.001 between *π* and gene density, for Finnish and Koryaks, respectively. The failure of Gossmann and colleagues^40^ to detect further correlations was attributed to the small variation of *N_e_* observed or to only having considered neutral diversity (synonymous changes). In another analysis of genome *N_e_* heterogeneity, Jiménez-Mena and colleagues^41^ also found significant variation in *N_e_* across the genome of cattle populations using the temporal *N_e_* estimation method, but this variation did not correlate with the recombination rate, the density of genes, or the presence of loci under artificial selection. This negative result was attributed to the assumption of large genomic windows in order to have a large enough number of markers in each of them, or to the fact that the temporal method of estimation of *N_e_* was only based on a single-generation interval.^41–42^

The correlations found between the different parameters analyzed with genomic data follow the expected trends (Fig. S2) and agree with previous estimations. For example, analyzing 100 kb windows of a Danish population, Lohmueller and colleagues^49^ found significant correlations between the recombination rate and the number of SNPs in the windows (*r* = 0.20), the SNP diversity (*r* = 0.11) and SNP MAF (*r* = 0.062). These correlations are comparable with those found in our study between *RR* and genome diversity parameters (Fig. S2).

The large reduction in *N_eπ_* with respect to *N_eLD_* in Figure 1 for low recombination rates confirms that linked selection can reduce very substantially neutral genetic diversity. This result may have an implication on Lewontin’s^50^ paradox, i.e. the observation that differences in genetic diversities at neutral sites for most eukaryotes and prokaryotes species fall within a range of an order of magnitude,^51–53^ whereas differences in the average census sizes between species differ in more than three orders of magnitude. This has been explained by alternative hypotheses. One of them suggests that species with large population sizes would generally have a high variation in reproductive success and would suffer from a higher impact from demographic changes.^54^ Another explanation is the possible constrain in genomic variation imposed by natural selection via fixation of beneficial mutations and elimination of deleterious ones in linked neutral variation.^55^ Theoretical developments showing that there is a nonlinear relationship between effective population size and census size, may contribute to explain this paradox.^27^ In this sense, because *N_eLD_* is not affected by selection, it would be expected to be closer to population size *N* than to *N_e_* from nucleotide diversity.

In summary, our results show that the estimates of historical effective size obtained from linkage disequilibrium between pairs of SNPs are not altered by selection, either positive or negative, and are not either strongly affected by the heterogeneity in recombination rate across the genome. Therefore, linkage disequilibrium *N_e_* reflects the true demographic changes in population size over generations. In contrast, other methods based on mutation and recombination, from which recent estimates of human demography have been obtained, can be strongly affected by selection.

## Methods

### Computer simulations

Genomic data of populations under different demographic and evolutionary scenarios were simulated with the software SLiM 3.^56^ Random mating populations of constant or changing size (ranging between *N* = 100 and 10,000) were run for up to 10,000 generations. Different demographic scenarios (constant population size, bottlenecking, exponential growth, etc.) were assumed. Genome sequences with a length of 100 Mb were considered where mutations occur at a rate between 10^-7^ and 10^-9^ mutations per nucleotide and generation, depending on the demographic scenario and simulation, with different recombination rates (ranging from 0.01 to 5 cM per Mb). For each locus, values of fitness of 1, 1 + *s*/2, and 1 + *s* were considered for the wild-type homozygote, the heterozygote, and the mutant homozygote, respectively. Fitness of an individual was assumed to be multiplicative across loci. Three mutation models were assumed. (1) A neutral model for all mutations (*s* = 0). (2) Background selection (BS), where 90% of mutations are neutral and 10% are deleterious with selection coefficient *s* = −0.02. (3) Selective Sweeps (SS), where 99% of mutations are neutral and 1% are assumed to be advantageous with effect *s* = 0.02. To investigate the heterogeneity of the genome in recombination rates, the simulated sequence was divided in 70 regions of equal length and the particular rate of recombination for each region was randomly chosen from the distribution of recombination rates observed in analogous genome windows of the human genome (Suppl. Figure S1). Each simulation was run for up to 100 replicates.

### Analysis of human genomes

Data comes from the genomic sequencing of samples from two human populations: 99 individuals from a Finnish population,^46^ with a total of about 9.4 million SNPs, and 16 individuals from a Koryaks’ population,^47^ with about 4.6 million SNPs. The Koryaks population data coordinates, in genome version hg18, were converted to hg19 using liftOver UCSC tool.^57^ In the process, 65.088 variants were not found and were excluded from the analyses. However, there is a large number of SNPs available in both populations, allowing the study of relatively small regions of the genome. Only autosomal chromosomes were taken into account. Genomic data was divided in 2 cM regions in which local *N_e_* and other genetic variables were estimated to investigate the correlations between one another. For an accurate estimation of linkage disequilibrium *N_e_*, only regions with more than 250,000 pairs of SNPs were considered. In addition, telomeric regions shorter than 2 cM and regions with very noisy *N_e_* outlier estimates (> 30,000) or negative ones were also excluded. Thus, following these criteria, 321 and 655 regions were excluded from the analyses of Finnish and Koryaks data, respectively, and the final number of regions analysed was 1,502 and 1,154, respectively.

### Estimation of *N_e_*

The software GONE was used to obtain historical estimates of linkage disequilibrium *N_e_* (*N_eLD_*) using all pairs of SNPs available from simulation data at distances between *c* = 0.5 and 0.001 Morgans (M) in samples of 100 individuals. The software MSMC^18^ and Relate^19^ were applied to the same data, except that only samples of four randomly sampled individuals were analysed with MSMC because of computation time restrictions with this software. MSMC version 2 (downloaded in December 2019) was used with the “fixedRecombination” flag, as recommended by the user’s guide. Since the software needs several chromosomes to be run, sets of 10 replicates were run and considered as chromosomes. Relate (downloaded in December 2019) was run without monomorphic SNPs, providing the mutation rate of the simulations, the number of haplotypes of the sample, a seed, 300 bins and a threshold value from 50 to 30 depending on the simulation scenario. It was run for each simulation replicate and the results averaged over replicates.

For the analysis of specific genomic regions with real data, *N_eLD_* estimates were directly obtained with equations S4b and S5 of the Supplemental Material of Santiago et al.^13^, which applies to the scenario of constant population size of diploid populations when the genetic phase of genotypic data is unknown. In this case, because SNPs in the regions are necessarily at relatively low genetic distances, pairs of SNPs at distances ranging between 1/50 and 1/100 M were considered in order to obtain at least 250,000 pairs per genomic region. The software Relate and MSMC were not used in these analyses, as they are assumed to apply only to historical estimates of *N_e_*.

### Estimation of other genomic variables

Recombination rates (*RR*; in cM/Mb) between all pairs of consecutive SNPs for each of the genomic regions were obtained from the human genetic map^34^ and averaged for each region. Estimates of the background selection statistic (*B*)^58^ were obtained for each site and averaged for each genomic region. A reduction in neutral diversity at a given genomic region is a function of the intensity of purifying selection and the rate of recombination, as the impact of selection on reducing diversity is higher in tight linkage regions.^22,26^ The *B* statistic measures the impact of background selection on nucleotide diversity. Thus, it fluctuates between one (no background selection affecting diversity) and zero (almost complete exhaustion of diversity as a result of background selection), with an average for the human autosomal genome of about 0.74 - 0.81.^58^

Average nucleotide diversity (*π*), polymorphism (*P*) and minor allele frequency (MAF) of SNPs were calculated for each genomic region. The number of Loss of Function (LoF) and missense variants in each genomic region were also obtained using data from the 0.3.1 version of the ExAC browser,^59^ downloaded on 14th October 2019. Only high confidence variants were taken into account. The gene density of each genomic region was obtained using the RefSeq database.^60^ When a gene spanned over different regions, we considered it to be in the region were its middle point was located. Only genes with a straightforward chromosome code were used (e.g. NC_000001.10 corresponding with chromosome 1).

## Acknowledgements

We thank Humberto Quesada for useful comments on the topic. This work was funded by Agencia Estatal de Investigación (AEI) (PID2020-114426GB-C21), Xunta de Galicia (GRC, ED431C 2020-05) and Centro singular de investigación de Galicia accreditation 2019-2022, and the European Union (European Regional Development Fund - ERDF), Fondos Feder “Unha maneira de facer Europa”. I.N. was funded by a predoctoral (FPU18/04642) grant from Ministerio de Ciencia, Innovación y Universidades (Spain).

## Declaration of interests

The authors declare no competing interests.

## Web resources

LiftOver UCSC tool, http://genome.ucsc.edu <u>RefSeq database</u>, https://ftp.ncbi.nlm.nih.gov/refseq/H_sapiens/annotation/GRCh37_latest/refseq_identifiers/GRCh37_latest_genomic.gff.gz

## Data availability

The authors affirm that all data necessary for confirming the conclusions of the article are present within the article, figures, and tables. The code generated during this study is available at github address […].

## Author Contributions

**Conceptualization:** Enrique Santiago, Armando Caballero.

**Formal analysis:** Irene Novo.

**Supervision:** Enrique Santiago, Armando Caballero.

**Writing – review & editing:** Armando Caballero, Irene Novo, Enrique Santiago.

## Notes

### Competing Interest Statement

The authors have declared no competing interest.

